# Modelling acceptable novelty transitions with emotional habituation: Effects of uncertainty and prediction error on preference changes

**DOI:** 10.1101/2020.08.27.269811

**Authors:** Takahiro Sekoguchi, Hideyoshi Yanagisawa

## Abstract

Previous mathematical models developed to optimize the degree of novelty in product design have represented novelty by emotional dimensions such as arousal (surprise) and valence (positivity and negativity). Formalizing arousal as Bayesian information gain and valence as a function of arousal based on Berlyne’s arousal potential theory, the model indicates that novelty becomes acceptable with repeated exposure and changes the preferences of the users. Hence, acceptable novelty transitions are important in the design of long-term product experience. We propose a mathematical model of acceptable novelty transitions with emotional habituation based on the emotional dimension model. We formalized valence as a function of information-theoretic free-energy and expressed free-energy as a function of three parameters: initial prediction error, initial uncertainty, and noise of sensory stimulus. To verify whether the transition speed of acceptable novelty depends on the initial uncertainty in our model, we analysed the responses of participants to historic artworks; we manipulated the uncertainty level by varying the obscurity of the presented pieces and the prediction error by rendering them in different artistic styles. We used the subjective reports of valence in response to the samples as measures of valence levels. The experimental results support our hypothesis.

## 1. Introduction

Novelty is an important aspect of attraction in product design. Whether the novelty of products is acceptable, however, depends on the preferences of the user. Raymond Loewy, a pioneer of industrial design, redefined the negotiation between novelty and acceptability as that between novelty attraction and the fear of the unknown; summarized by the acronym MAYA (most advanced, yet acceptable), this concept underscores the need for innovative designs to be widely accepted in society (Loewy, 2002).

Berlyne’s arousal potential theory suggests that pleasant feelings can be maximized when an appropriate arousal potential in the range between the familiar and novel and between the simple and complex is reached (Berlyne, 1967, 1970, 1971). However, the arousal potential that maximizes positive hedonic response may differ according to an individual’s knowledge or experience(Giacalone, Duerlund, Bøegh-Petersen, Bredie, & Frøst, 2014; Silvia, 2005).

Many studies have proposed emotional dimension models based on two-dimensions: arousal and valence (Lang, 1995; Russell, 1980). Arousal indicates the intensity of the arousal state. For example, emotions such as surprise are associated with high arousal and bored state is classified as low arousal. Valence indicates positive or negative feeling. Informed by information theory and the Bayes model, We previously developed a mathematical model of emotional dimensions associated with novelty that considered personal characteristics (Yanagisawa, Kawamata, & Ueda, 2019). The model formalized arousal (primary emotional dimension) as a Kullback–Leibler(KL) divergence(Kullback & Leibler, 1951) from Bayesian posterior to prior, or information gain. We formulated the information gain as a function of prediction error, uncertainty, and external noise using Gaussian prior and posterior. Prediction error and uncertainty represented the level of novelty and personal characteristics, respectively. The function model was experimentally supported using the event-related potential (ERP) P300 of human participants as an index of arousal. Furthermore, they formalized valence as a function of arousal based on Berlyne’s arousal potential theory. We recently found that information theoretic free energy (or surprisal) represents arousal potential and includes the information gain(or Bayesian surprise)(Yanagisawa, 2020).

Individuals adapt to novelty by experiencing it repeatedly: i.e., habituation, or the response extinction to the repeated presentation of stimuli (Houtveen, Rietveld, Schoutrop, Spiering, & Brosschot, 2001; Ludden, Schifferstein, & Hekkert, 2012). Habituation to novelty and changes in emotions associated with habituation are important in the design of long-term product experience. Having formulated a mathematical model of habituation based on the emotional dimension model (Yanagisawa et al., 2019), We previously modelled habituation to novelty as a decrement of information gain through Bayesian update and found that the initial prediction error and uncertainty affect habituation(Sekoguchi, 2019). Our model was experimentally supported using the ERP. Lévy et al. (Lévy, MacRae, & Köster, 2006) demonstrated that the range of novelty and complexity that maximizes pleasant feelings increased with repeated exposure to stimuli. This phenomenon is caused by changes in valence resulting from the increased habituation to novelty with the repeated exposure to stimuli because the habituation to novelty is accompanied by changes in valence. However, mathematical models that can explain habituation-induced changes in valence have received little attention in the literature.

This study sought to develop a mathematical model to explain and predict transitions in valence caused by habituation to novelty. We extents our models of emotion dimensions (Sekoguchi, 2019; Yanagisawa et al., 2019) to consider the valence transitions in novelty habituation. We formulate the framework for a mathematical model of valence transitions using the free-energy(Friston, Kilner, & Harrison, 2006) as arousal potential(Yanagisawa, 2020), and analysed the effects of initial prediction error and initial uncertainty on valence transitions with the model. Finally, we validated the model predictions in an experiment that presented stimuli of varying uncertainty levels and prediction errors to adult participants.

## 2. Modelling acceptable novelty transitions

### a. Emotional arousal model

In this section, we review our emotional arousal model in terms of habituation to novelty(Sekoguchi, 2019; Yanagisawa et al., 2019). Novelty was defined as the amount of information acquired when an event is experienced. The information content given by an event *x* termed self-information is described as (*x*) ln *p(x*), where *p*(*x*) is the probability of *x ∈* ***R***. Self-information is consistent with its uncertainty before experiencing it. An expected value of self-information (i.e. Shannon’s entropy(Shannon, Weaver, Blahut, & Hajek, 1949)) of the distribution represents the uncertainty of the prior expectation and is described as following:

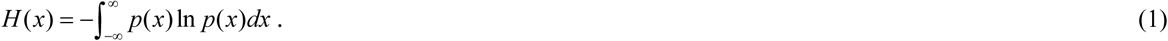

Considering the state transition across experiencing an event, we defined belief distributions before and after the transition as *prior p* and *posterior q*, respectively. The Shannon’s entropy decreases from prior to posterior. This decrease is proportional to the information acquired from the event, which we defined as *information gain* [5]. Information gain is described as the following:

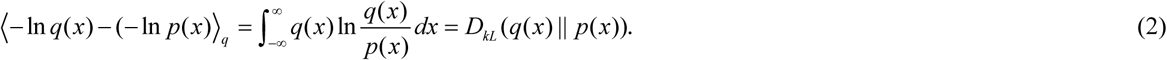

The expression *D*_*KL*_(*p*∥ *q*) is the KL divergence of *q* from *p* (Kullback & Leibler, 1951). The information gain represents the human surprise that attracts human attention(Itti & Baldi, 2009; Yanagisawa et al., 2019).

Let *π(θ*) be a *prior* of a parameter *θ* that one estimates. The parameter can be regarded as a cause of sensory input. After one obtains continuous data *x*∈***R*** (e.g. sensory input) by experiencing an event, the prior is updated to the posterior *π (θ*|*x*) using Bayes’ theorem:

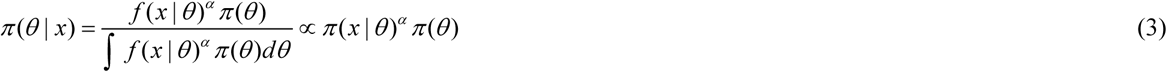

where *f*(*x*|*θ*) is a likelihood function of *θ* when one obtains data *x*, and *a* is a parameter that adjusts the amount of the update: i.e., the *learning rate*. The Bayesian posterior represents distributions of perception of *θ* after obtaining data *x* (Yanagisawa, 2016).

The posterior when the same events are experienced *k* times, *π*_*k*_(*θ*|*x*), is considered to be updated to the prior when the same data are obtained *k*+1 times, *π*_*k+1*_(*θ*). Assuming that the likelihood functions are i.i.d, the order of data collection is irrelevant. Thus, the posterior when *k*^*th*^ data obtains is the following equation:

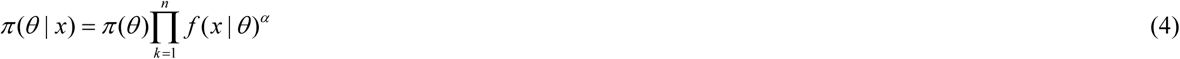

The posterior *π*_*k*_(*θ*|*x*) is proportional to the product of the initial prior and each likelihood function. Assume that one obtains *n* samples of the same data *x* and encodes them as a Gaussian distribution *N*(*µ, s*_*l*_) with a flat prior. Further assume that one has a non-flat prior of *µ* that follows a Gaussian distribution *N*(*µ*_*p*_, *s*_*p*_). Using the formula (9), the prior is updated to a Gaussian distribution *N*(*µ*_*post*_, *s*_*post*_) when obtaining the same data *n* times, where

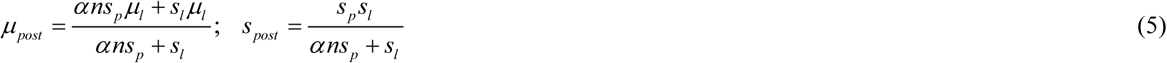

In these equations, *µ*_*l*_ is the mean of the likelihood. From formula (5), the information gain from the prior to the Bayesian posterior (or Bayesian surprise(Itti & Baldi, 2009)) when obtaining the same data *n* times *G*_*n*_ can be derived as follows:

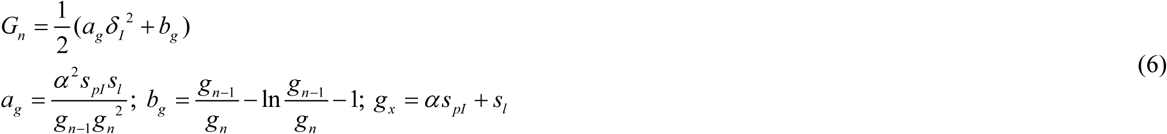

where *δ*_*l*_ |*µ*_*p*_ *− µ*_*l*_| is termed *initial prediction error*, representing the difference between expectations and data that one initially obtains; *s*_*pl*_ is termed *initial uncertainty* because it is proportional to the information entropy of the *initial prior* (e.g. before obtaining data *x*); and *s*_*l*_ is termed *external noise*, representing the variance of the data. From formula (6), the information gain can be formulated as a function of the three parameters: initial prediction error *δ*_*l*_, initial uncertainty *s*_*pl*_, and external noise *s*_*l*_.

### b. Free-energy principle

We modelled valence using the free-energy principle, the theory that uniformly explains recognition, perception, and action human living (Friston et al., 2006). Free-energy is defined as the energy required for livings to process information in their brain. This theory assumes that adaptive systems such as human brains encode a probabilistic model of the causes of their sensations and must minimize the amount of free-energy to resist a natural tendency to disorder.

When a biological agent perceives external state *π (θ*), it is sent to the agent’s brain as sensations, and the brain generates the recognition density *q*(*θ*). The brain has a probabilistic generative model *π (θ, x*) that projects the relationship between sensory samples and their causes. Free-energy is defined as follows:

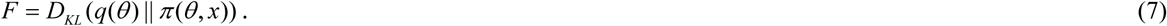

The Bayesian theorem provides two alternative expressions for free-energy. The first defines free-energy as the KL divergence between the recognition density and posterior and self-information of sensations called *sensory surprise*. In this case, recognition density must approach the posterior to minimize free-energy because the lower limit of free-energy is sensory surprise.

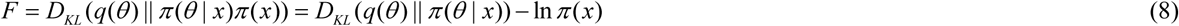

The second defines free-energy as the KL divergence between the recognition density and prior and likelihood averaged over recognition density, called *inverse accuracy*.

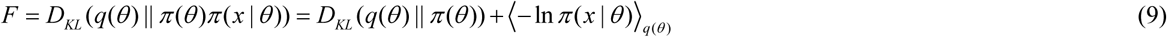

When the recognition density approaches the posterior to minimize free-energy, free-energy can be decomposed as a summation of a KL divergence between the prior and posterior and negative-log-likelihood averaged over posterior. The first term is *Bayesian surprise*(Itti & Baldi, 2009) that correspond to information gain used as an index of arousal(Yanagisawa et al., 2019). The second term is the inverse accuracy, meaning the uncertainty in recognition of the parameter *θ* remaining after experiencing an event caused by noisy sensations. For example, when one drinks a mixture of orange and grapefruit juice, they cannot be sure whether the drink was orange or grapefruit. This is because we consider the inverse accuracy as an index of uncertainty. The second term can be interpreted as perceived complexity of stimuli under recognition of their causes (Yanagisawa, 2020). In summary, minimized free-energy contains both arousal and uncertainty.

### c. Free-energy as arousal potential

Berlyne assumed the existence of a range that maximizes the pleasure between the novel and familiar and between simple and complex stimuli [2]. Further, he assumed that these hedonic qualities of stimuli arise from separate biological incentivization systems, the reward and aversion systems, which can be represented by a sigmoid function [2]. The joint operation of these two systems generates an inverted U-shaped curve called a Wundt curve. The valence of a stimulus changes from neutral to positive as the arousal potential increases but shifts from positive to negative once the arousal potential passes the peak positive valence.

Novel and complex stimuli yield large arousal potential and require a large amount of energy to process; hence, people tend to avoid them. On the other hand, while simple and familiar stimuli are associated with peace of mind, they do not inform adaptations to environmental changes. People prefer moderately novel and complex stimuli that save energy while providing new information. Free-energy is compatible with the energy required to process information about novel or complex stimuli(or uncertain causes). Free-energy can, therefore, be considered as an index of arousal potential.

### d. Modelling valence as a function of free-energy

Assuming prior distribution and a likelihood function that follows a Gaussian distribution, free-energy when experiencing an event *n* times is obtained from the following formula (10):

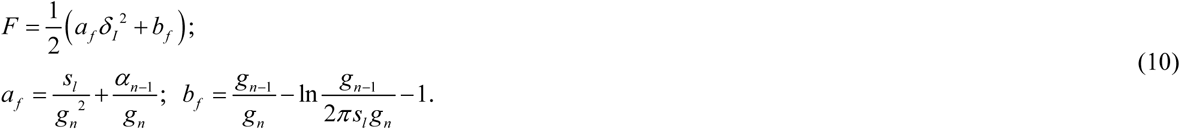

Valence can be expressed as the sum of the reward and aversion systems (Yanagisawa et al., 2019), both of which are represented by the sigmoid function and are a function of free-energy.

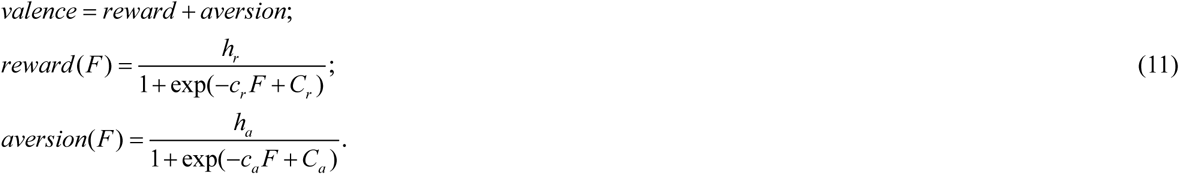

In (11), *C*_*r*_, *h*_*r*_, and *c*_*r*_ represent the thresholds of the information gain that activate the reward systems, the maxima of positive valence levels, and the gradients, respectively; the corresponding values in the aversion system are *C*_*a*_, *h*_*a*_, and *c*_*a*_, respectively.

## 3. Simulation and analysis

We analysed how free-energy changes as the number of updates increases. Equation (12) is equal to the partial derivatives of free-energy with respect to the number of updates – i.e., the decrease rate of free-energy.

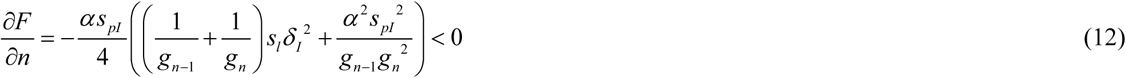

Equation (12) implies free-energy decreases as the number of updates increases. We analysed how the initial prediction error and initial uncertainty affect the decrease of free-energy. The partial derivatives of the decrease rate of free-energy with respect to the initial uncertainty are less than zero:

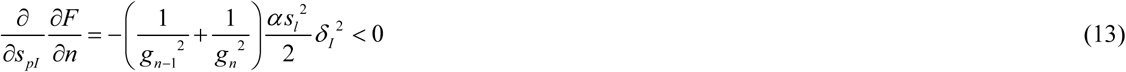

Equation (13) implies that the decrease rate of free-energy increases as the initial uncertainty increases, and the difference of the decrease rate of free-energy between different initial uncertainties rises as the initial prediction error increases. Figure 1 shows the relationship between the number of updates and the decrease rate of free-energy (formulated in (13)) for different initial uncertainties. The greater the initial uncertainty, the more rapidly free-energy decreases. Figure 2 shows that the difference in the decrease rate of free-energy between different levels of initial uncertainty becomes more apparent as the initial prediction error rises.

**Figure 1:**
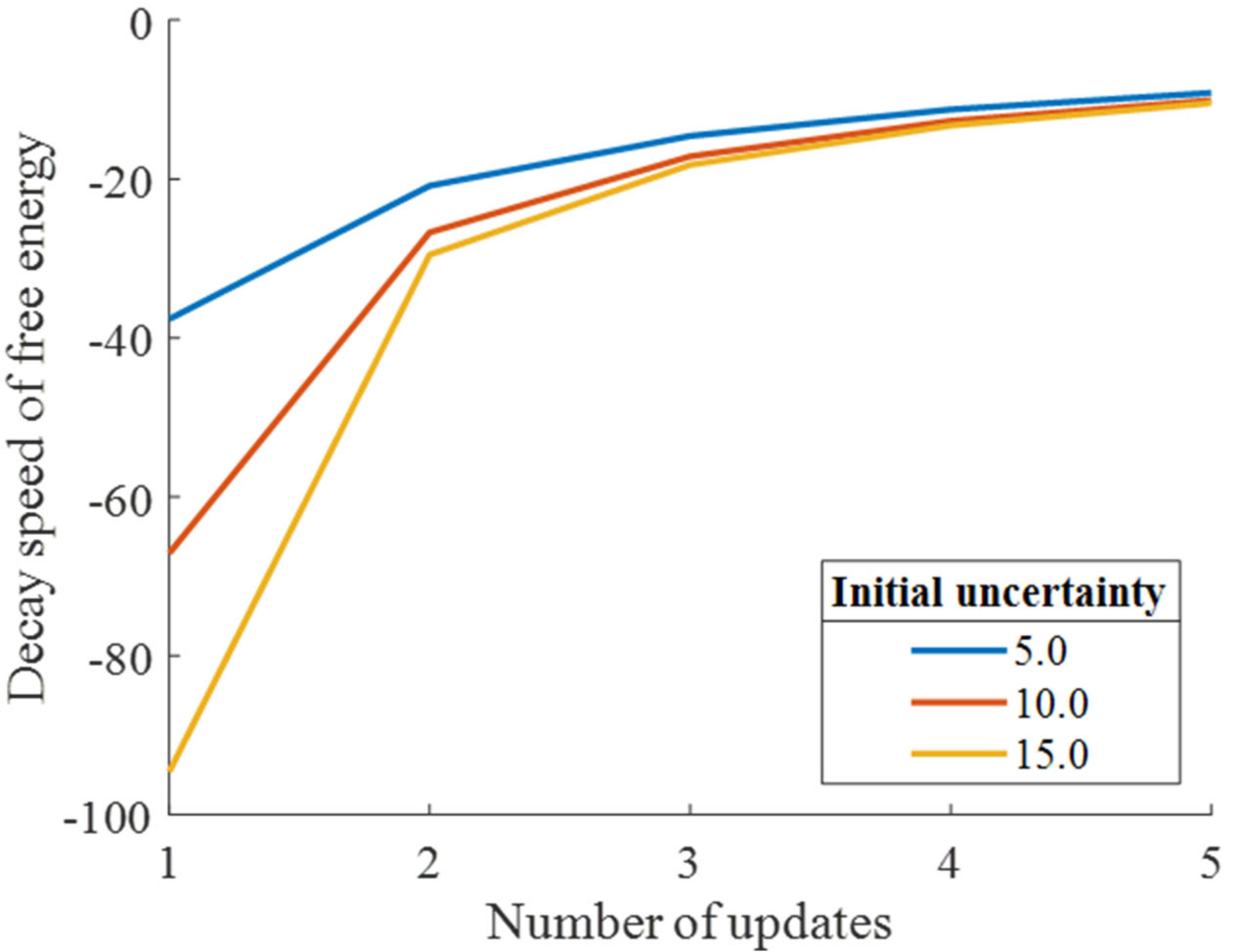
Decrease rate of free-energy as a function of the number of updates for different initial uncertainties. The decrease rate of free-energy rises as the initial uncertainty increases for any number of updates.

**Figure 2:**
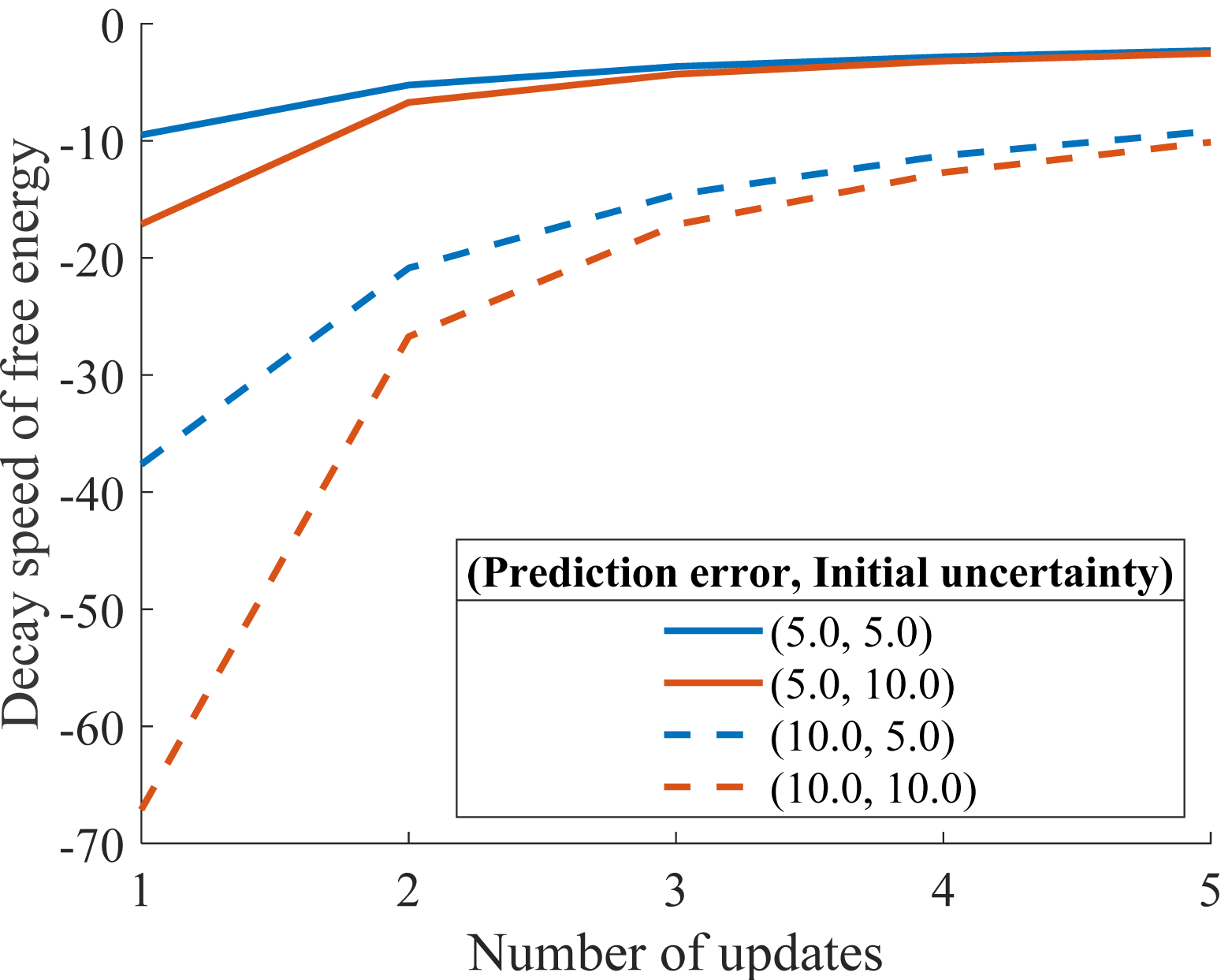
Decrease rate of free-energy as a function of the number of updates for different combinations of initial uncertainties and initial prediction errors. The difference in the decrease rate of free-energy between different levels of initial uncertainty becomes more apparent as the initial prediction error increases.

The more the free-energy decreases, the greater valence shifts along the Wundt curve from large to small arousal. For example, as shown in Figure 3, a stimulus that people feel to be unpleasant upon first encounter is likely to become less unpleasant as the experience is repeated. This phenomenon is known as the mere exposure effect. This model provides for the increased rate of free-energy decrease as the initial uncertainty increases: i.e., the mere exposure effect tends to occur among individuals with less expectation for an event.

**Figure 3:**
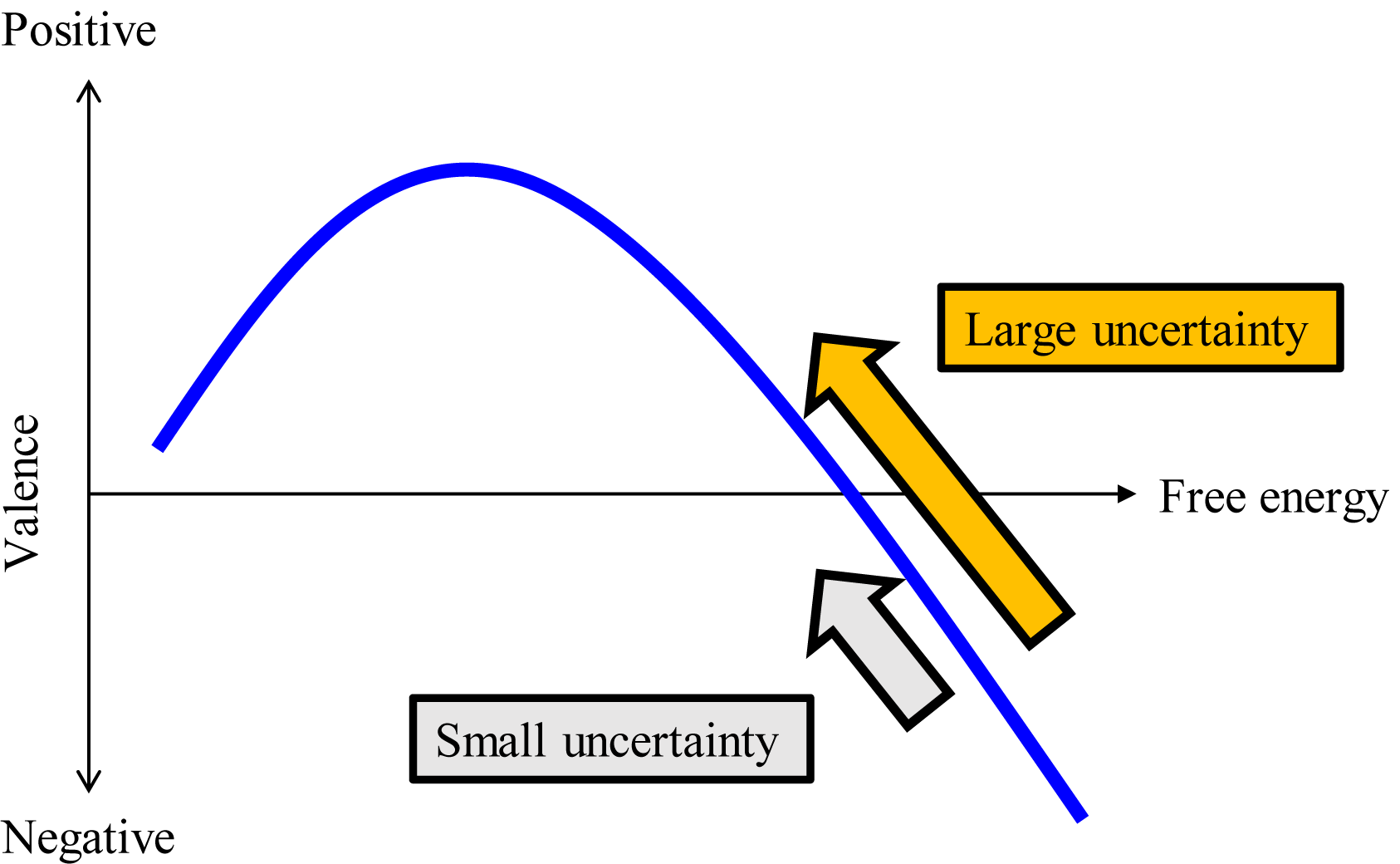
Comparison of the speed of the valence shift on the Wundt curve between different uncertainties. The greater the decrease in free-energy, the greater the transition of the valence change along Wundt curve from large to small arousal. The larger the initial uncertainty, the more rapidly the free-energy decreases. Hence, the larger the initial uncertainty, the greater the acceptability of novelty transitions.

## 4. Experiment

We conducted an experiment to validate our model’s ability to predict the valence transitions discussed in section 3. The experiment of the model prediction tested the hypothesis that the greater the initial uncertainty and the greater the initial prediction error, the sooner the valence transition.

### a. Participants

The present study enrolled 108 adults (age range: 20-49, 54 male, 54 female). The experimental protocol was approved by the Ethics Committee of the Graduate School of Engineering at the University of Tokyo. In accordance with the principles of the Declaration of Helsinki, all participants provided written informed consent prior to their participation in this study. The participants could stop the experiment sessions at their convenience.

### b. Stimuli

We used paintings as visual stimuli to obtain data on the emotional valence of the participants’’ responses to different initial prediction errors and uncertainties. We controlled the initial prediction error by manipulating the art style of a picture. Specifically, mixed an original painting with another that is representative of a different style by using an art-style transfer model and a convolutional neural network(Gatys, Ecker, & Bethge, 2016). We thus rendered original works in two different styles (Figure 4).

**Figure 4:**
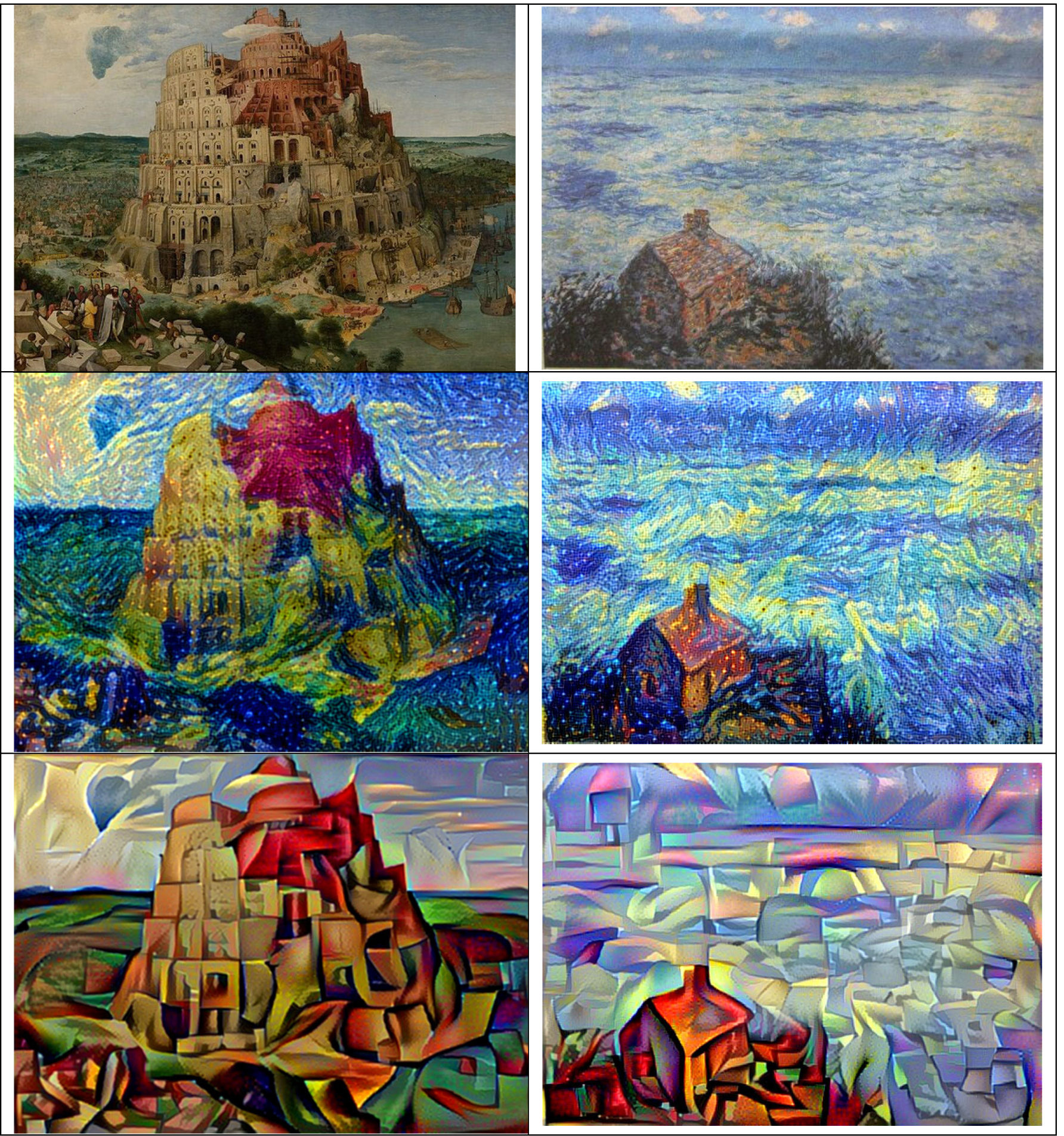
Examples of experimental stimuli. The upper pictures are original samples. The middle pictures are rendered as post-impressionist (small prediction error), and the lower are rendered as cubist (large prediction error). We varied the familiarity of the samples to expand the scope of the data: left panel, “The Tower of Babel” by Bruegel, more recognizable; right panel, “View Over the Sea” by Monet, less recognizable.

Table 1 shows all the combinations of the original pictures and those with which they were mixed to render the originals in different styles. We selected eight paintings by Monet and Bruegel as representative examples of Impressionism or the Renaissance, respectively, to vary the familiarity of the samples. We mixed the eight pictures with those produced by van Gogh or Picasso as representative examples of Post-Impressionism and Cubism, respectively. We assumed that when the original pictures were mixed with the starkly contrasting style of Cubism, the generated image would yield a greater prediction error from the original than when the original painting was mixed with the less contrasting style of Post-Impressionism; i.e., we assumed a Bruegel-Picasso combination would prompt more perceived incongruity than would a Bruegel-van Gogh painting (Figure 4). All pictures were obtained from Wikipedia Commons and were thus free of copyright. At the beginning of the experiment, we asked the participants whether he or she had seen the eight original pictures. We used their responses to inform the level of uncertainty for each of the original pictures: a recognized picture generates more concrete expectations concerning the rendition of the pictures than one that has never been seen before. A detailed description of the procedure is provided in the next section.

**Table 1:** Combinations of original pictures with those of different styles

| No. | Original pictures<br>(Impressionism or<br>Renaissance) | Pictures mixed to the original picture |  |
| --- | --- | --- | --- |
|  |  | Post-Impressionism<br>(small prediction error) | Cubism<br>(large prediction error) |
| 1 | Bruegel<br>“The Tower of Babel” | Gogh “The starry night” | Picasso<br>“The Weeping Woman” |
| 2 | Monet<br>“La Grenouillere” | Gogh “The starry night” | Picasso<br>“The Weeping Woman” |
| 3 | Monet<br>“La Gare Saint-lazare” | Gogh “The starry night” | Picasso<br>“The Weeping Woman” |
| 4 | Monet<br>“La Manneporte” | Gogh “The starry night” | Picasso<br>“The Weeping Woman” |
| 5 | Monet<br>“View Over the Sea” | Gogh “The starry night” | Picasso<br>“The Weeping Woman” |
| 6 | Monet<br>“Twilight, Venice” | Gogh “The starry night” | Picasso<br>“The Weeping Woman” |
| 7 | Monet<br>“The Japanese Footbridge” | Gogh “Enclosed Field with<br>Ploughman” | Picasso<br>“The Weeping Woman” |
| 8 | Monet<br>“Impression, Sunrise” | Gogh “The starry night” | Picasso<br>“The Weeping Woman” |

### c. Procedure

The experiment consisted of two sections. In the first section, we presented the eight unedited paintings to the participants and asked them whether they were familiar with any of them. We used pictures that participants had seen before for the high initial uncertainty stimulus and one that the participant had never seen before for the low initial uncertainty stimulus. In the second section, we presented either the picture of the post-impressionist or cubist version of painting along with the picture of the original painting. (see Table 1 for the pairs). The participants were divided into two groups according to the order in which the modified pictures were presented: in group A, the first four randomly presented pairs featured the original painting rendered in a post-impressionist style, while the second four featured the cubist paintings; this order was reversed in group B. Each set was presented in a random order to the participants four times. In each trial, after presenting the pair of pictures to the participants, they were asked: “How do you feel about modified picture relative to the original picture?” Participants could select from seven pre-prepared responses that corresponded to the valence states A-D along the Wundt curve and the midpoints between them (Figure 5): level A indicates less arousal and boring; B, moderate arousal and interesting; C, large arousal and acceptable; D, extreme arousal and unacceptable. Hence, the seven responses – represented by the range of numbers between 1 and 7 – included A, between A and B, B, between B and C, C, between C and D, and D, respectively. We assumed that valence shifts from D to A through the repeated exposure to the same paintings.

**Figure 5:**
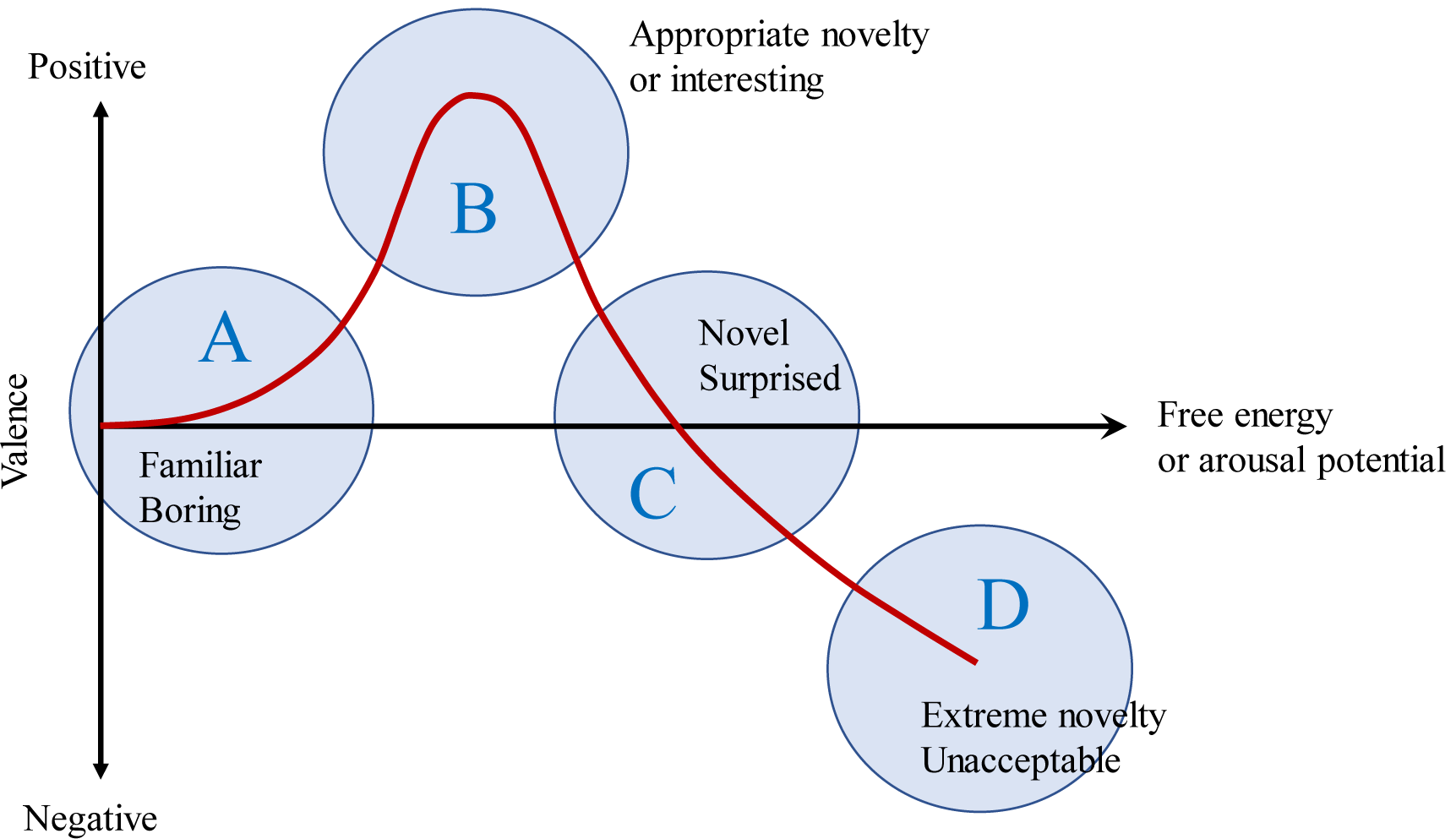
Four levels of valence (A-D) the participants referenced in responding to the experimental stimuli.

## 5. Results

Of the artworks rendered in the post-impressionist style, 46 had never been seen before (small uncertainty) in groups A or B, and 186 had been seen before (large uncertainty). Of those rendered in the cubist style, 40 had never been seen before, and 181 had been seen before.

Figure 6 showed how the state values of the participants’ responses, as reflected by a position on the Wundt curve, changed from the first to fourth exposure for the two styles (i.e., prediction error). The x-axes of both figures represented the numbers of repeated exposures to a stimulus. The y-axes represent the changes in the state values of the participants’ responses (ranging from 1 to 7) from the first exposure; e.g., if the state changed from D to the midpoint between C and D, the amount of change was registered as one. The state value changes obtained in response to samples featuring the same initial uncertainty and prediction error were averaged. By comparing the amount of change in the state value between the first and fourth exposure, we investigated whether participants had ever appreciated samples or not affected the change of their answer through four times of exposure regardless of their preference for the samples at the first exposure. Post-impressionist samples (small prediction error) did not prompt any significant differences in the rates of the state-value change according to whether the participants had seen the samples or not (Figure 6(a)); by contrast, the cubist samples (large prediction error) did (Figure 6(b)). This finding indicates that the aesthetic preference of the participants who had never seen the samples shifted along the Wundt curve towards less arousal. The resulting difference in the state change between different levels of initial uncertainties corresponds to the model prediction shown in Figure 2.

**Figure 6:**
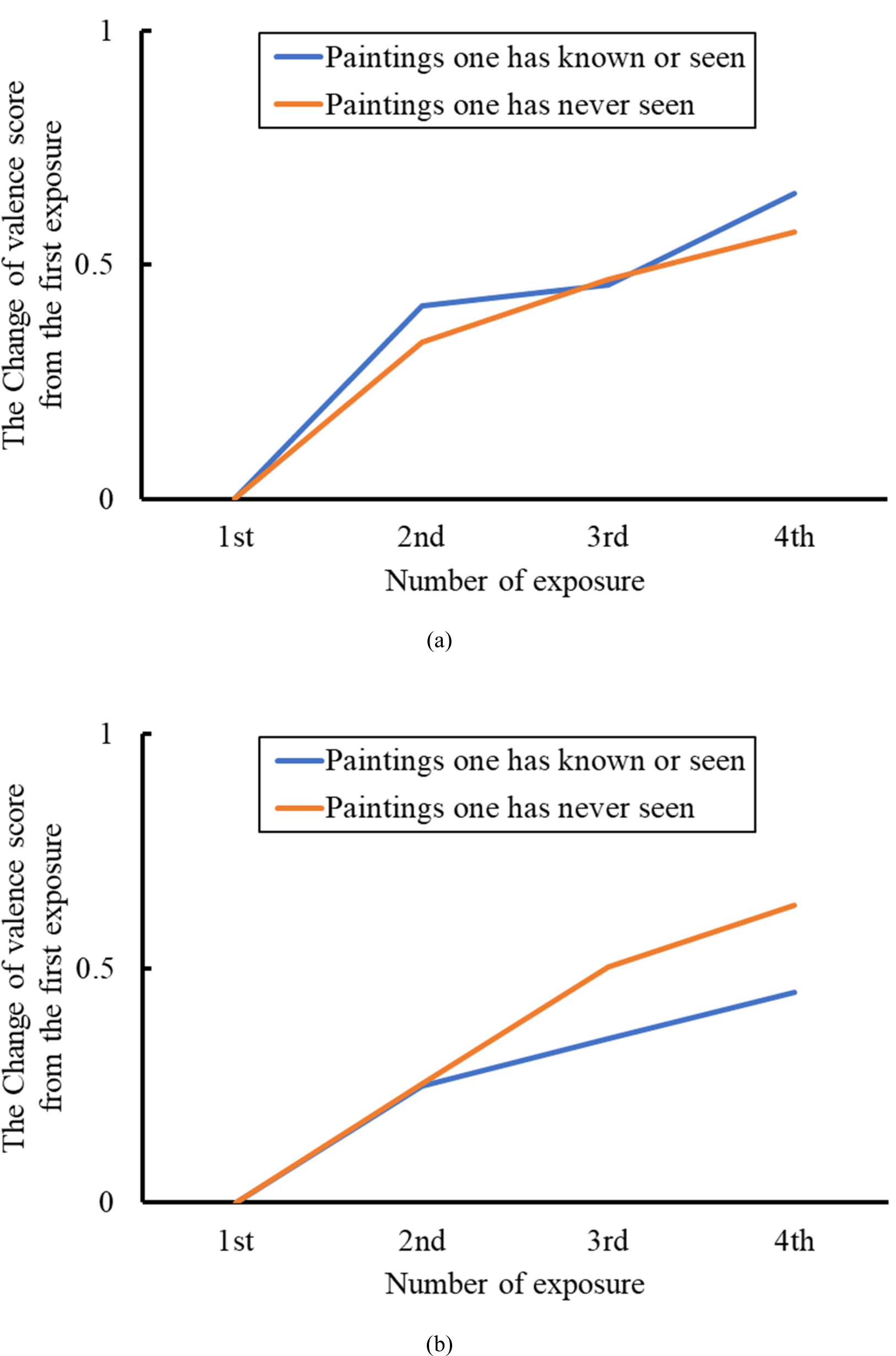
State transitions along the Wundt curve due to repeated exposure of the same picture rendered in different styles: (a) post-impressionist and (b) cubist. The colour of the line indicates whether the participants had seen or known of the picture before the experiment.

## 6. Discussion and Conclusion

We assumed that free-energy, i.e., the energy living things use to process even-related information, represents arousal potential, an index of novelty and complexity in Berlyne’s arousal potential theory. We suggested that the minimized free-energy used to adapt to an environment can be deconstructed into information gain and inverse accuracy: the indices of arousal and inverse accuracy, respectively. Free-energy is generated when experiencing a novel event as a function of the parameters of the initial prediction error, initial uncertainty, and external noise when both prior and likelihood function are assumed to follow Gaussian distributions. We analysed how the initial prediction error(difference between initial prior mean and likelihood peak) and initial uncertainty(variance of initial prior) affects free-energy when the same events are experienced repeatedly. The model predictions show that free-energy decreases with the repeated experience of the same event and that the lower rates of free-energy decrease further as the initial uncertainty increases. Furthermore, the greater the initial prediction error, the greater the difference in the decreased rate of free-energy between different levels of initial uncertainty. These results indicate that knowledge about a novel stimulus is negatively associated with the preference for stimulus change: e.g., our model predicts non-specialists of art are more likely than experts to shift dynamically among works when appreciating complex arts.

To validate our model prediction, we conducted an experiment that manipulated the styles of historic paintings. Specifically, we rendered Renaissance and impressionist artworks in the styles of later periods. We chose artworks that differed in their anticipated familiarity among the participants to vary initial uncertainty levels. While we did not find significant differences in the valence change between initial uncertainties when we rendered the original artworks in the post-impressionist style, participants who had never seen the original works changed their preferences more dynamically once the paintings were rendered in the cubist style. The initial prediction error indicates the deviation of an event from an expectation informed by knowledge or experience: in the present case, the difference between the presentation of an artwork and how a participant expected to see it. Our results suggest that the initial prediction error affect how preferences concerning works change with repeated exposure.

The agreement between our model prediction and experimental results suggests that free-energy obtained from a novel and complex event can be considered an index of Berlyne’s arousal potential. We formulated free-energy as a function of initial prediction error and initial uncertainty. As both initial uncertainty and initial prediction error depend on knowledge and experience, our proposed model explains how preferences change through experience and the factors that cause this change. Our study assumed emotional habituation to novelty and acceptable novelty transition as a temporary phenomenon. The only situation tested in this experiment was the appreciation of several pictures in a short time. In the real world, experience is shaped over long periods, and the impressions of novel events change dynamically. Hence, future research on the interaction between novelty and experience should consider longer periods.

## Acknowledgments

We thank Prof. Tamotsu Murakami and the members of the Design Engineering Laboratory at the University of Tokyo for supporting this project.

## Ethical Statement

This study was approved by the Research Ethics Committee of the University of Tokyo, Graduate School of Engineering (Approval number KE19-107).

## Data Accessibility

The stimuli we used in this experiment and the experimental results are available in supplemental material.

## Authors’ Contributions

HY supervised the study. TS and HY formalized and analysed the mathematical model. TS and HY designed the experiment. TS analysed the experimental data. TS and HY drafted the manuscript. All authors revised the manuscript.

